# Rotational dynamics in motor cortex are consistent with a feedback controller

**DOI:** 10.1101/2020.11.17.387043

**Authors:** Hari Teja Kalidindi, Kevin P. Cross, Timothy P. Lillicrap, Mohsen Omrani, Egidio Falotico, Philip N. Sabes, Stephen H. Scott

## Abstract

Recent studies hypothesize that motor cortical (MC) dynamics are generated largely through its recurrent connections based on observations that MC activity exhibits rotational structure. However, behavioural and neurophysiological studies suggest that MC behaves like a feedback controller where continuous sensory feedback and interactions with other brain areas contribute substantially to MC processing. We investigated these apparently conflicting theories by building recurrent neural networks that controlled a model arm and received sensory feedback about the limb. Networks were trained to counteract perturbations to the limb and to reach towards spatial targets. Network activities and sensory feedback signals to the network exhibited rotational structure even when the recurrent connections were removed. Furthermore, neural recordings in monkeys performing similar tasks also exhibited rotational structure not only in MC but also in somatosensory cortex. Our results argue that rotational structure may reflect dynamics throughout voluntary motor circuits involved in online control of motor actions.

**Highlights:**

- Neural networks with sensory feedback generate rotational dynamics during simulated posture and reaching tasks
- Rotational dynamics are observed even without recurrent connections in the network
- Similar dynamics are observed not only in motor cortex, but also in somatosensory cortex of non-huma n primates as well as sensory feedback signals
- Results highlight rotational dynamics may reflect internal dynamics, external inputs or any combination of the two.

## Introduction

Motor cortex (MC) plays an important role in our ability to make goal-directed motor actions such as to reach and grasp objects of interest in the environment. A key approach to explore MC’s contribution to movement has been to record the patterns of neural activity during tasks such as reaching. In the last part of the 20^th^ century, research emphasized the representation of movement parameters by cortical networks (Fetz, 1992; Scott, 2008; Vyas et al., 2020). This approach assumed that activity of individual neurons or at the population level could be directly related to explicit features of motor action such as movement speed or muscle activity patterns.

However, there has been a recent transition towards interpreting neural processing using dynamical systems techniques (Machens et al., 2010; Michaels et al., 2016; Pandarinath et al., 2018b, 2018a; Remington et al., 2018; Russo et al., 2018; Sauerbrei et al., 2020; Shenoy et al., 2013; Suresh et al., 2020). Churchland et al., (2012) recorded from MC while monkeys performed goal-directed reaches and fit the population activity to an autonomous dynamical system where future activity was predicted based solely on the past population activity in MC. They found this relationship could account for a significant amount of the neural activity and revealed rotational dynamics that could provide a basis set for generating muscle activity patterns. However, these rotational dynamics are absent in supplementary motor cortex suggesting that they are not trivial properties of cortical processing (Lara et al., 2018).

This view of MC as a pattern generator during reaching was further bolstered by recurrent neural network models (RNN) (Hennequin et al., 2014; Michaels et al., 2016; Sussillo et al., 2015). RNNs trained to generate patterns of muscle activity while constrained to generate simple dynamics also displayed rotational dynamics that resembled MC activity (Sussillo et al., 2015). Importantly, these networks only received external inputs that were stationary with the exception of a non-selective GO cue to initiate the pattern generation. Thus, activity was generated solely by the connections between neurons and online feedback about the generated muscle patterns was not necessary after training. Collectively, these results have led to the interpretation that the function of MC is to generate patterns of muscle activity and that this real-time process is done largely autonomously from other brain structures.

Another class of dynamical systems is also commonly used in motor control to interpret the behavioural aspects of motor actions. Specifically, a growing body of literature has highlighted how optimal feedback control (OFC) can capture how we move and interact in the world (Franklin and Wolpert, 2011; Scott, 2004, 2016; Shadmehr and Krakauer, 2008; Todorov and Jordan, 2002). OFC highlights the importance of feedback processes, both external sensory feedback (e.g. proprioception and vision) as well as internal feedback from efference copies, for generating motor commands for movement. A large number of studies inspired by OFC highlight how humans are capable of generating fast, goal-directed motor corrections (Cluff and Scott, 2015; Cross et al., 2019; Dimitriou et al., 2012; Kurtzer et al., 2008; Nashed et al., 2014; Scott, 2016) even for very small disturbances (Crevecoeur et al., 2012) and OFC can capture features of unperturbed movements (Knill et al., 2011; Lillicrap and Scott, 2013; Liu and Todorov, 2007; Nashed et al., 2012; Todorov and Jordan, 2002; Trommershäuser et al., 2005). Further studies highlight how feedback responses to a mechanical disturbance are distributed throughout somatosensory, parietal, frontal and cerebellar motor circuits in ~20ms and display goal-directed responses in as little as 60ms (Chapman et al., 1984; Conrad et al., 1975; Evarts and Tanji, 1976; Herter et al., 2009; Lemon, 1979; Omrani et al., 2016; Phillips et al., 1971; Pruszynski et al., 2011, 2014; Strick, 1983; Wolpaw, 1980). This interpretation of motor control emphasizes that the objective of the motor system is to attain the behavioural goal and this requires feedback processed by a distributed network. Further, MC is generally viewed as part of the control policy that uses information on the system state to generate muscle activity patterns to attain the behavioural goal.

These two views of MC, one as an autonomous dynamical system and the other as a flexible feedback controller, appear to conflict on how to interpret the role of MC and its interactions with the rest of the motor circuits involved in goal-directed motor actions. This apparent conflict seems to hinge on the observation that the rotational dynamics observed in MC can be generated through purely local recurrent connections. However, it is unclear if a feedback control network would also exhibit similar rotational dynamics and whether these dynamics are exclusively in MC or also in other brain regions such as somatosensory cortex. We investigated this question by first developing a multi-layer RNN that controlled and received sensory feedback from a two-segment limb. The network was trained to counter disturbances to the limb and perform reaching movements. After training, rotational dynamics were observed in the network activities as well as in sensory feedback from the limb, but not in muscle activity. Critically, rotational dynamics could also be generated with or without recurrent connections in the trained networks. Monkeys trained in a similar task exhibited rotational dynamics in MC and also in somatosensory and posterior parietal cortices including during reaching where sensory feedback is not required a priori. Taken together, these results illustrate rotational dynamics can be observed across frontoparietal networks and can be generated by intrinsic dynamics in MC and/or through dynamics of the entire motor system.

## Results

### RNN exhibit rotational dynamics in the activities and sensory feedback signals during posture task

Rotational dynamics in MC have been interpreted as a signature of an autonomous dynamical system (Churchland et al., 2012; Pandarinath et al., 2018a; Shenoy et al., 2013). In contrast, rotational dynamics appear to be absent in systems that are dominated by external inputs, such as muscle activity driven by neural inputs (Churchland et al., 2012), or MC activity during grasping driven by sensory inputs (Suresh et al., 2020). Here, we examined the dynamics of a network performing a posture perturbation task, where the network had to respond to sensory feedback about the periphery to generate an appropriate motor correction (Cross et al., 2020; Heming et al., 2019; Omrani et al., 2014, 2016; Pruszynski et al., 2014). Sensory input plays an important role for correctly performing the task and thus the hypothesis is that rotational dynamics should be absent in the network.

We built an artificial neural network that controlled a two-link model of the upper limb (Figure 1). Previous neural network models (Hennequin et al., 2014; Michaels et al., 2016; Sussillo et al., 2015) focused on network activities (*r*) that evolved according to 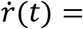 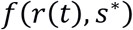 where *f*[∙]is a nonlinear function and *s*\* is vector of static inputs about the GO cue and the current target. Here, we generated a model where network activities also incorporated delayed (Δ) continuous sensory feedback about the limb (s(t-Δ)) and thus activities evolved according to 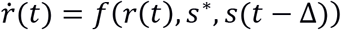. The neural network contained an input layer that had recurrent connections between neurons and received delayed (50ms) sensory feedback about the limb state (i.e. joint position, velocity, muscle activities). This layer projected to an output layer that also had recurrent connections between neurons. The output layer directly controlled the activities of six muscles (two sets of monoarticular muscles at the shoulder and elbow joints and two biarticular muscles) that generated limb movements. The network was trained to perform a posture perturbation task where the goal was to keep the limb within a specified target location, while countering randomly applied loads to the limb. We optimized the network by minimizing a cost function that penalizes the kinematic error between the target location and current limb position over the duration of the task.

**Figure 1.**
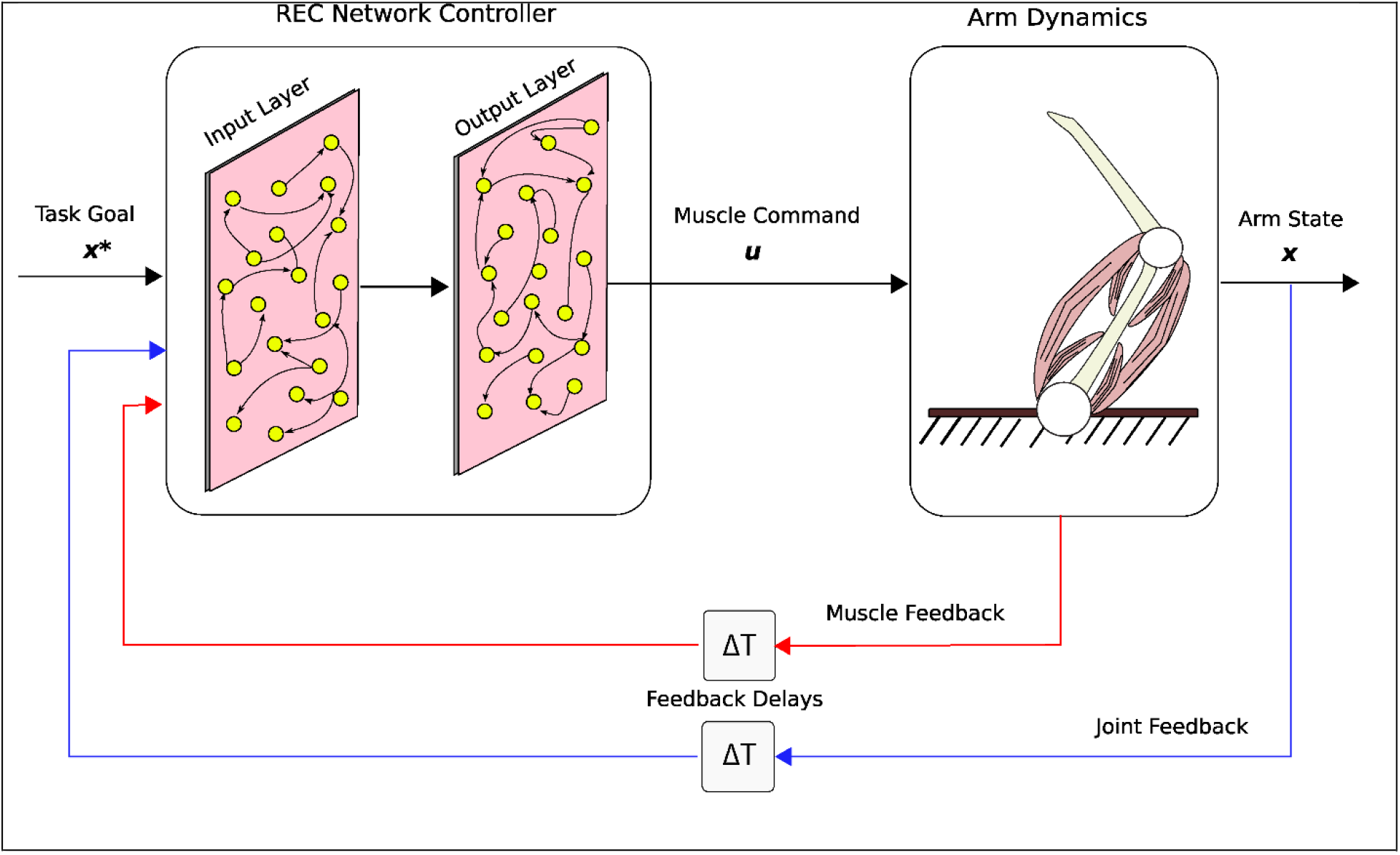
Simulation setup. Schematic of the two-link model of the arm and the neural network. The arm had two joints mimicking the shoulder and elbow (Arm Dynamics: joints are white circles) and was actuated using 6 muscles (pink banded structures). Muscle activity was generated by the neural network (Muscle Command). The network was composed of two layers (Input and Output layers) with recurrent connections between units within each layer. The network received delayed (ΔT) sensory feedback from the limb in the form of joint angles and velocities (Joint Feedback, blue line), and muscle activities (Muscle Feedback, red line). Delays were set to 50ms to match physiological delays. The network also received input about the desired location of the limb (Task Goal).

After optimization we applied loads that displaced the limb by ~3cm. The network generated corrections to the displacements with the hand reversing direction within 300-400ms from the time of the applied load (Figure 2A-C). The network also maintained steady-state motor output for the remainder of the trial to counter the applied loads. Figure 2D shows the activity of the shoulder extensor muscle aligned to the load onset. An increase in muscle activity started 50ms after the applied load, consistent with the delay in sensory feedback from the limb. Muscle activity peaked at ~200ms after the applied load and stabilized to a steady state within ~750ms. Figures 2E and F show the activity of two example neurons from the output layer of the network.

**Figure 2.**
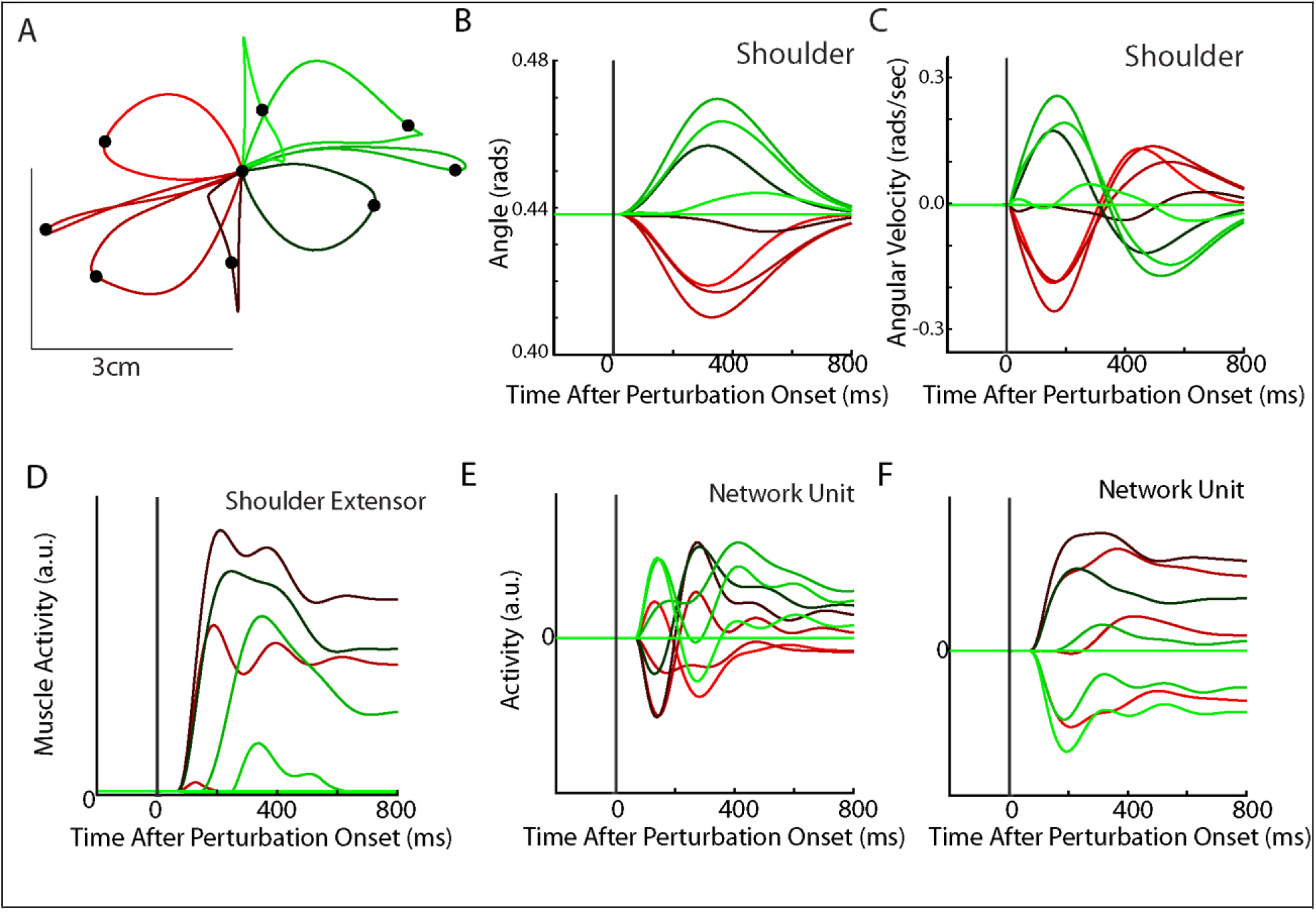
Posture perturbation task performed by neural network. A. Hand paths when mechanical loads were applied to the model’s arm. Due to the anisotropy in the biomechanics the trajectories across the different loads are asymmetric. Black dots denote the hand’s location 300ms after the load onset. B-C) Shoulder angle and angular velocity aligned to the load onset. D) Activity of the shoulder extensor aligned to load onset. E-F) The activities of two example units from the output layer of the network. The colors in A-F correspond to different directions of load.

We examined the population dynamics of the output layer of the network by applying jPCA analysis (Churchland et al., 2012). Briefly, jPCA constructs a multi-dimensional matrix (*X*(*t*), dimensions n × ct) which is composed of each unit’s (n) activity patterns across time (t) and condition (c) (e.g. load combination or reach target). The matrix is reduced (*X*_*Red*_)to a 6 × ct dimensional matrix using principal component analysis (PCA) to examine the dynamics exhibited by the dominant signals. This matrix is then fit to a constrained dynamical system 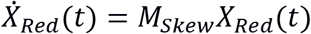 where 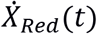 is the temporal derivative of *X*_*Red*_(*t*), and *M*_*Skew*_ is the weight matrix constrained to be skew symmetric. The skew-symmetric constraint ensures that only rotational dynamics are fit to the population activity and *M*_*Skew*_ can then be decomposed into a set of three jPC planes.

We found the top-2 jPC planes exhibited clear rotational dynamics with rotation frequencies of 2.0Hz and 0.7Hz (Figure 3A, left and middle panels). Combined, these two planes captured 60% of the variance of the output-layer activities. In contrast, the third jPC plane exhibited a more expansion-like property (Figure 3A, right) and captured 38% of the variance. Examining the goodness of fit (R^2^) to the constrained dynamical system provides a measure of how well the activities in the network activities are approximated by rotational dynamics. We compared our results to a null distribution that tested whether the rotational structure was an emergent property of the population activity or simply reflected known properties of single-neuron responses (i.e. broad tuning for loads, smooth time-varying activity patterns, shared patterns of activity across neurons). We used tensor maximum entropy (TME, Elsayed and Cunningham, 2017) to generate surrogate datasets that were constrained to have the same covariances as the observed data and applied the same jPCA analysis to the datasets. We found the constrained dynamical system had an R^2^ of 0.55 and was significantly greater than expected from the null distributions (Figure 3B left; TME: median R^2^=0.27, p=0.001). Further, when we did not constrain the weight matrix to be skew-symmetric (i.e. unconstrained dynamical system, M_Best_), we found an increase in the R^2^ to 0.83 that was also significant (Figure 3B right; median R^2^=0.49, p<0.001). The ratio between the R^2^ for the constrained and unconstrained fits was 0.66 indicating that the majority of the output layer’s dynamics displayed rotational dynamics.

**Figure 3.**
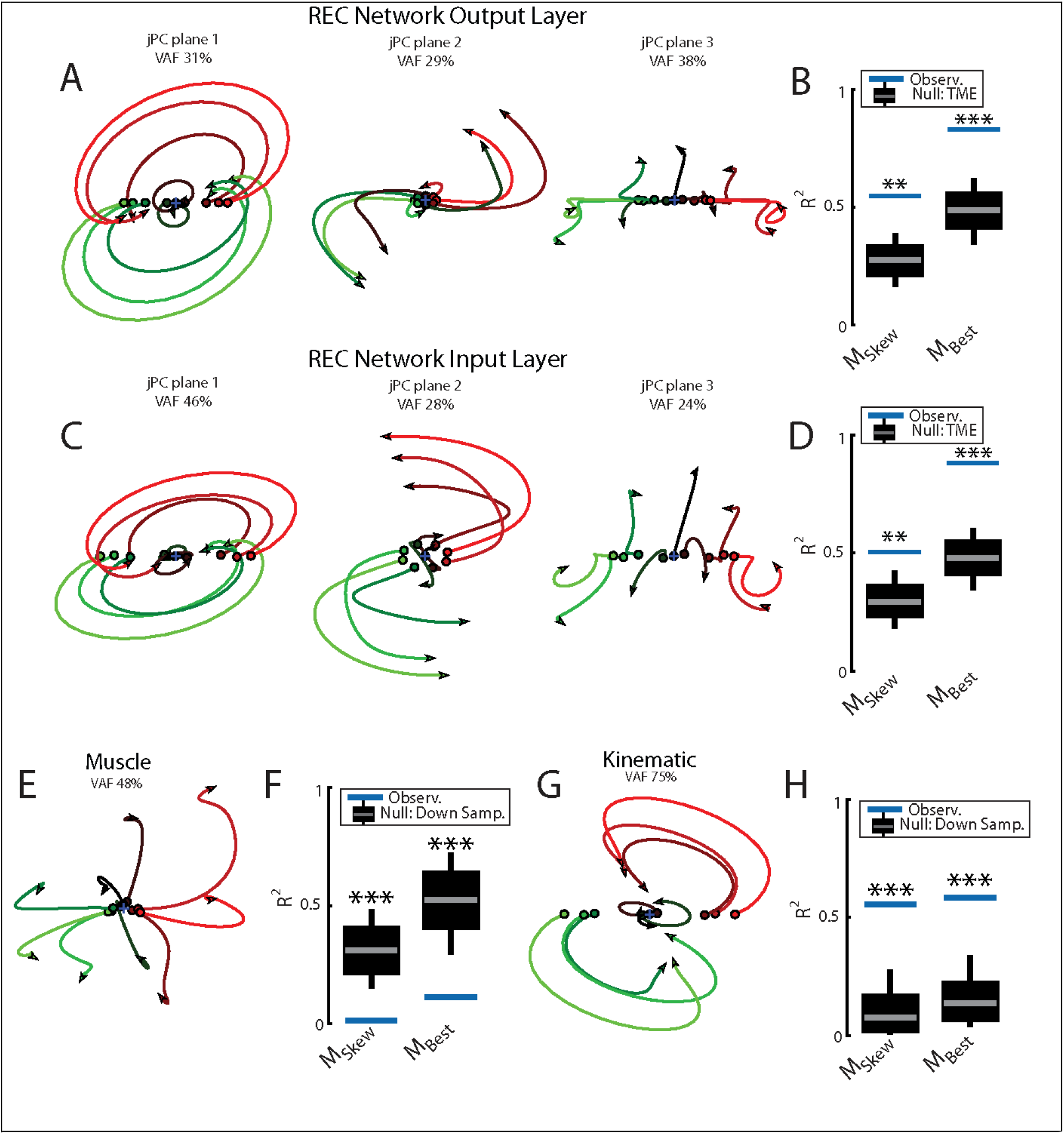
Population dynamics of the network during posture. A) The top-3 jPC planes from the activity in the output layer of the network. Dynamics were computed from 70ms to 370ms after the load onset. Different colours denote different load directions. VAF =variance accounted for. B) The goodness of fit (black horizontal line) of the network activity to the constrained (M_Skew_ left) and unconstrained (M_Best_ right) dynamical systems. Null distributions were computed using tensor maximum entropy (TME). Grey bars denote the median, the boxes denote the interquartile ranges and the whiskers denote the 10^th^ and 90^th^ percentiles. C-D) Same as A-B) except for the input layer of the network. E-F) and G-H) Same as A-B) except for the muscle activities and kinematic inputs into the network, respectively. Null distributions were computed from the down-sampled neural activity for F and H.

Next, we examined if rotational dynamics were present in the input layer of the network which directly receives sensory feedback. Similar to the output layer, we observed rotational dynamics in the top-2 jPC planes with frequencies of 1.8Hz and 0.95Hz (Figure 3C). Combined, these two planes captured 74% of the variance of the input-layer activity. The fit to a constrained dynamical system had an R^2^ = 0.51 (Figure 3D left) and was also significantly greater than the null distributions (median R^2^=0.29, p<0.01). When fit with an unconstrained dynamical system, we also found an increase in the R^2^ to 0.88 that was significant (Figure 3D right; median R^2^=0.48, p < 0.001). Thus, rotational dynamics are present in the input layer that directly received sensory feedback as well the output layer that formed the muscle signals.

Next, we explored if rotational dynamics were present in the motor outputs (i.e. muscle activities) and sensory inputs (i.e. muscle activities and joint kinematics) of the network. We applied jPCA analysis to the muscle activities and did not observe clear rotations in any of the jPC planes (Figure 3E). We found the muscle activities were poorly fit to the constrained (Figure 3F; R^2^ = 0.01) and unconstrained dynamical systems (R^2^ = 0.11). One explanation for this lower fit quality is that muscle activity has substantially fewer signals (6) than the network activities (500). We tested this by down-sampling neural units to match the number of muscles. Note, we did not compute a null distribution using TME as we found hypothesis testing using TME was unreliable when the number of signals were small (<30). We found the goodness of fits for muscle activities were significantly smaller than the down-sampled neural activities (Figure 3F, constrained p<0.001; unconstrained p=0.002) indicating that the down-sampled neural activity exhibited greater dynamical properties than muscle activity.

Next, we applied jPCA analysis to the kinematic signals (angle and angular velocity of the joints). We observed clear rotational dynamics in the top jPC plane (Figure 3G) with a rotational frequency of 1.3Hz. We found the constrained and unconstrained dynamical systems had an R^2^ = 0.56 and 0.59, respectively, which were significantly larger than the null distributions (Figure 3H; down sampled neural population: constrained and unconstrained p<0.001).

These results indicate kinematic signals exhibit substantial rotational dynamics; however, their rotational frequencies are lower than observed in the output layer activities. Here we asked whether these higher frequencies could be explained by combining all available sensory feedback (i.e. muscle and kinematics). We fit a linear model that decoded the output layer’s activity in each jPC plane using the sensory feedback signals composed of kinematic and muscle signals. We found the predicted activities were highly similar to the output layer activities (R^2^=0.99) with virtually identical frequencies of rotation (Figure Supplementary 1A). This indicates sensory feedback provided rich signals that could exhibit rotational dynamics identical to the network’s dynamics.

### Motor and somatosensory cortex exhibit rotational dynamics while monkeys performed posture perturbation task

Next, we examined if rotational dynamics exist in MC activity. We trained five monkeys to perform a similar posture perturbation task. The limb kinematics were qualitatively similar to the network with limb displacements of ~3cm and hand reversal starting in 300-400ms (Figure 4A-C). Muscle activity tended to be multi-phasic within the first 500ms after the applied load and reached a steady state within 800ms (Figure 4D). We also examined data from two previously collected monkeys performing a similar task using an endpoint manipulandum (data from Chowdhury et al., 2020). These monkeys also exhibited fast corrective movements to the load applied to the manipulandum (Figure S2A-C).

**Figure 4.**
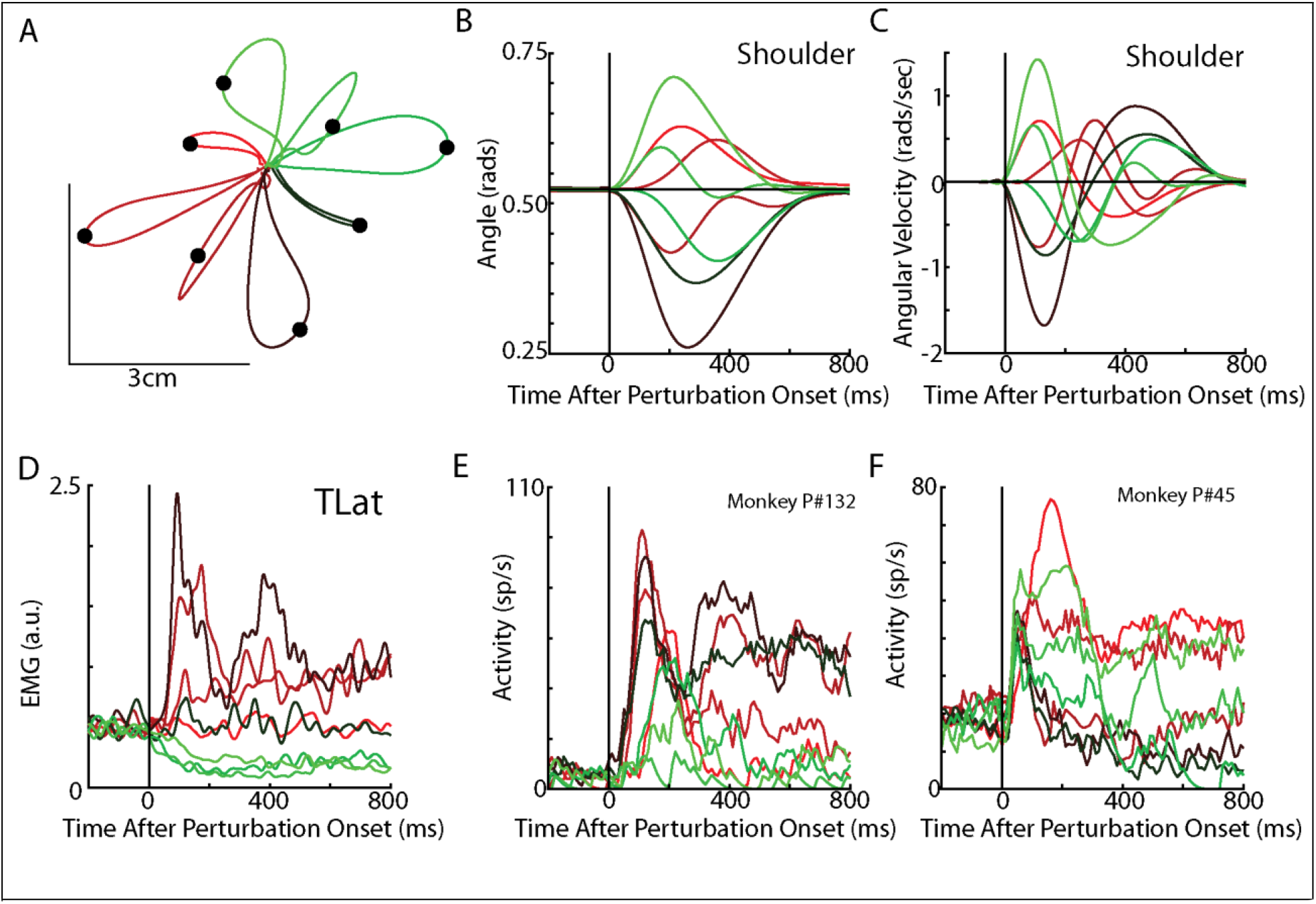
Posture perturbation task performed by monkeys. A) Hand paths for Monkey P when mechanical loads were applied to its arm. B-C) Shoulder angle and angular velocity aligned to the onset of the mechanical loads. D) Recording from the lateral head of the triceps (elbow extensor) during the posture perturbation task. E-F) Example neurons from motor cortex aligned to perturbation onset.

Neural activities were recorded using single electrodes (Monkeys P, A, X) and chronic multi-electrode arrays (Monkeys Pu, M, H and C). We observed motor cortex (MC) responses tended to peak in <200ms after the applied load and also exhibited steady-state activity (Figure 4E-F).

We pooled MC neurons across monkeys and then applied jPCA analysis. We found clear rotational dynamics in the top-2 jPC planes with frequencies of 1.3Hz and 1.1Hz for the first and second planes, respectively (Figure 5A). These planes also captured 63% of the variance from the neural population. In the third plane, we observed expansion-like dynamics similar to the third plane of the neural network (data not shown, 12% of variance). When we examined the fit qualities, we found the constrained and unconstrained dynamical systems had significant fits with an R^2^ of 0.41 (p<0.001) and 0.50 (p<0.001), respectively (Figure 5B blue lines, “Group Pop.”). Similar results were found when we applied jPCA for each monkey. For Monkeys P, A, X and Pu we found population activities exhibited rotational dynamics in the top-2 jPC planes (Figure S3A-D, rotation frequency range: plane 1=2.4-1.6Hz, plane 2=1.4-1.2Hz). Significant fits were found for the constrained (Figure 5B; mean across monkeys R^2^= 0.45, p<0.01) and unconstrained dynamical systems (mean R^2^= 0.56, p<0.05). However, for Monkey M we observed less rotational structure and more tangled trajectories in the top-2 jPC planes (Figure S3E). Fits for the constrained and unconstrained dynamical systems were still significant (constrained: p=0.003, unconstrained: p=0.002) but notably lower than for the other monkeys (constrained R^2^=0.21, unconstrained R^2^=0.32).

**Figure 5.**
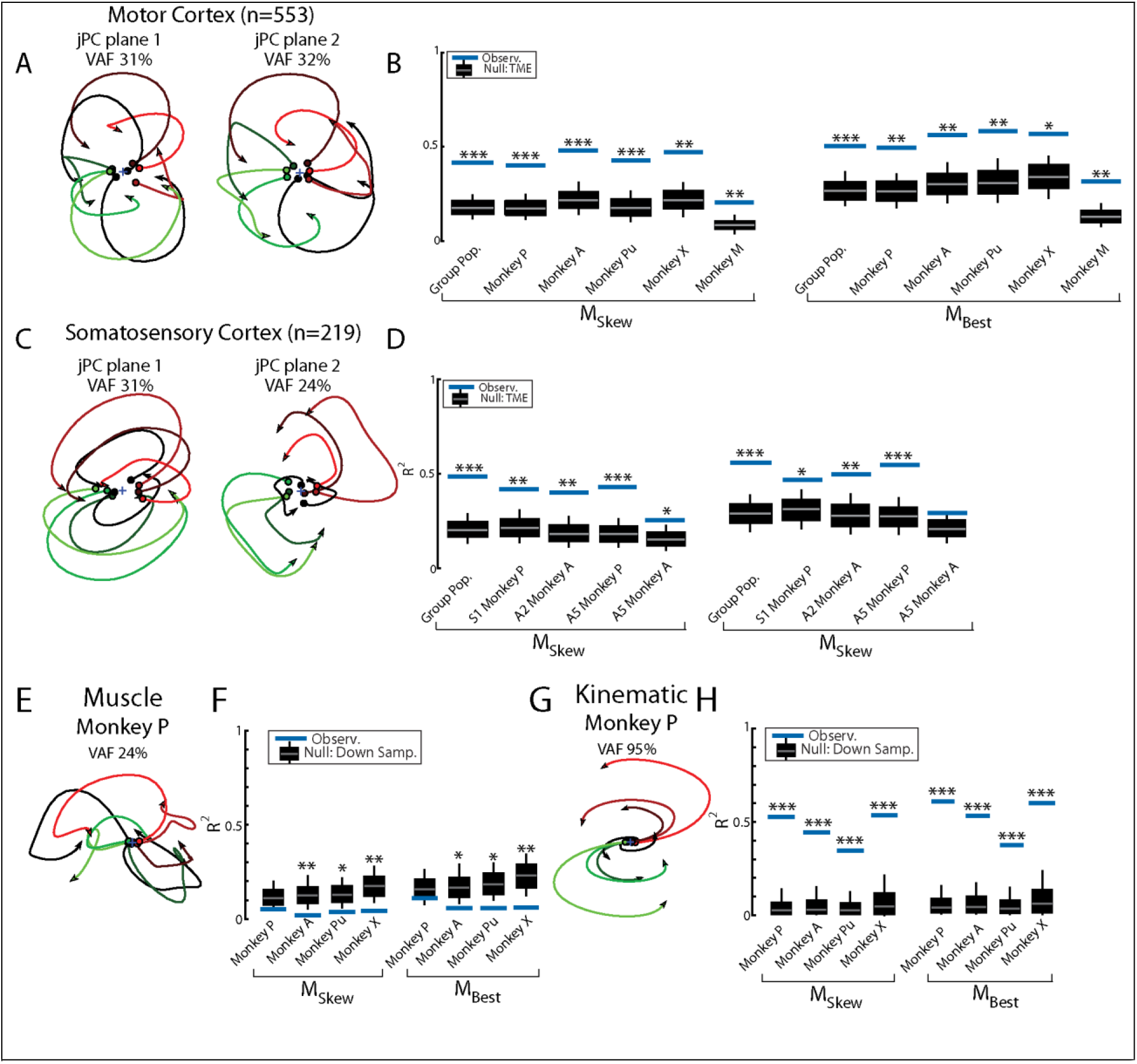
Population dynamics across motor and somatosensory cortex. A) The top-2 jPC planes from activity recorded in motor cortex pooled across all monkeys. B) Goodness of fits to the constrained (M_Skew_ left) and unconstrained (M_Best_ right) dynamical systems for motor cortex activity for the pooled activity across monkeys (Group Pop.) and for each individual monkey. Null distributions were computed using tensor maximum entropy (TME). C-D) Same as A-B) for somatosensory recordings. E) The top jPC plane from muscle activity from Monkey P. F) Goodness of fits to the muscle activity for the constrained and unconstrained dynamical systems for each monkey. G-H) Same as E-F) for kinematic signals. B,D, F, H) Grey bars denote the medians, the boxes denote the interquartile ranges and the whiskers denote the 10^th^ and 90^th^ percentiles. * p<0.05, **p<0.01, ***p<0.001.

We also examined the population dynamics in cortical areas associated with sensory processing (areas S1, A2 and A5). When neurons were pooled across monkeys, we observed clear rotational dynamics in the top-2 jPC planes with rotational frequencies of 1.7Hz and 1.1Hz (Figure 5C). Significant fits were found for the constrained (Figure 5D; R^2^=0.49, p<0.001) and unconstrained (R^2^=0.56, p<0.001) dynamical systems that were comparable to MC. Similar results were found when we applied jPCA for each monkey and cortical area separately (Figure 5D, S1D-E, S4).

Next, we examined the dynamics of the muscle activities and kinematic signals. We observed no rotational dynamics in the muscle activities for any of the monkeys (Figure 5E). We found the fits for the constrained and unconstrained dynamical systems were poor (Monkey P/A/Pu/X: constrained: R^2^=0.05/0.02/0.04/0.04, unconstrained: R^2^=0.11/0.06/0.06/0.06) and were significantly worse than the down-sampled neural activity (probability values plotted in Figure 5F). In contrast, for the joint kinematics we observed clear rotational dynamics with a rotation frequency of 1.3±0.1Hz (across monkeys mean and SD; Figure 5G, Figure S2F). We found the fits for the constrained and unconstrained dynamical systems were good (constrained: R^2^=0.45±0.03, unconstrained: R^2^=0.50±0.04) and significantly better than the down-sampled neural activity (probability values plotted in Figure 5H and Figure S2G). Lastly, for each monkey, we also decoded M1’s activity in each jPC plane using the joint kinematics and muscle activity and found the decoded activity was similar to M1’s activity (Figure S5).

### RNN exhibit rotational dynamics in the activities and sensory feedback signals during delayed reach task

Rotational dynamics were first described in MC during a delayed reaching task and inspired the interpretation of MC as an autonomous dynamical system (Churchland et al., 2012; Hennequin et al., 2014; Michaels et al., 2016; Sussillo et al., 2015). We explored if our network also exhibited similar rotational dynamics by training it on a delayed center-out reaching task. The plant dynamics and network architecture were the same as the posture task. However, the network was trained to maintain the limb at the starting location while a target was presented (“delay period”). Following a variable time delay, a ‘GO’ cue was provided requiring the network to move the limb to the target location within ~500ms.

After optimization, the REC network was able to generate limb reaches towards radially located targets at displacements of 2cm and 5cm from the initial location (Figure 6A). Reaches had bell-shaped velocity profiles, that peaked roughly during the middle of the movement (Figure 6B-C). Figure 6D shows the activity of the shoulder extensor muscle during reaches to different target locations. Figure 6E-F show the diverse temporal profiles exhibited by units in the output layer of the network. The unit in Figure 6E has a stable response during the delay period when the target was present. After the ‘GO’ signal, the unit exhibits oscillatory activity with a change in the unit’s preferred direction. The unit in Figure 6F largely maintains its preferred direction during the delay and movement periods.

**Figure 6.**
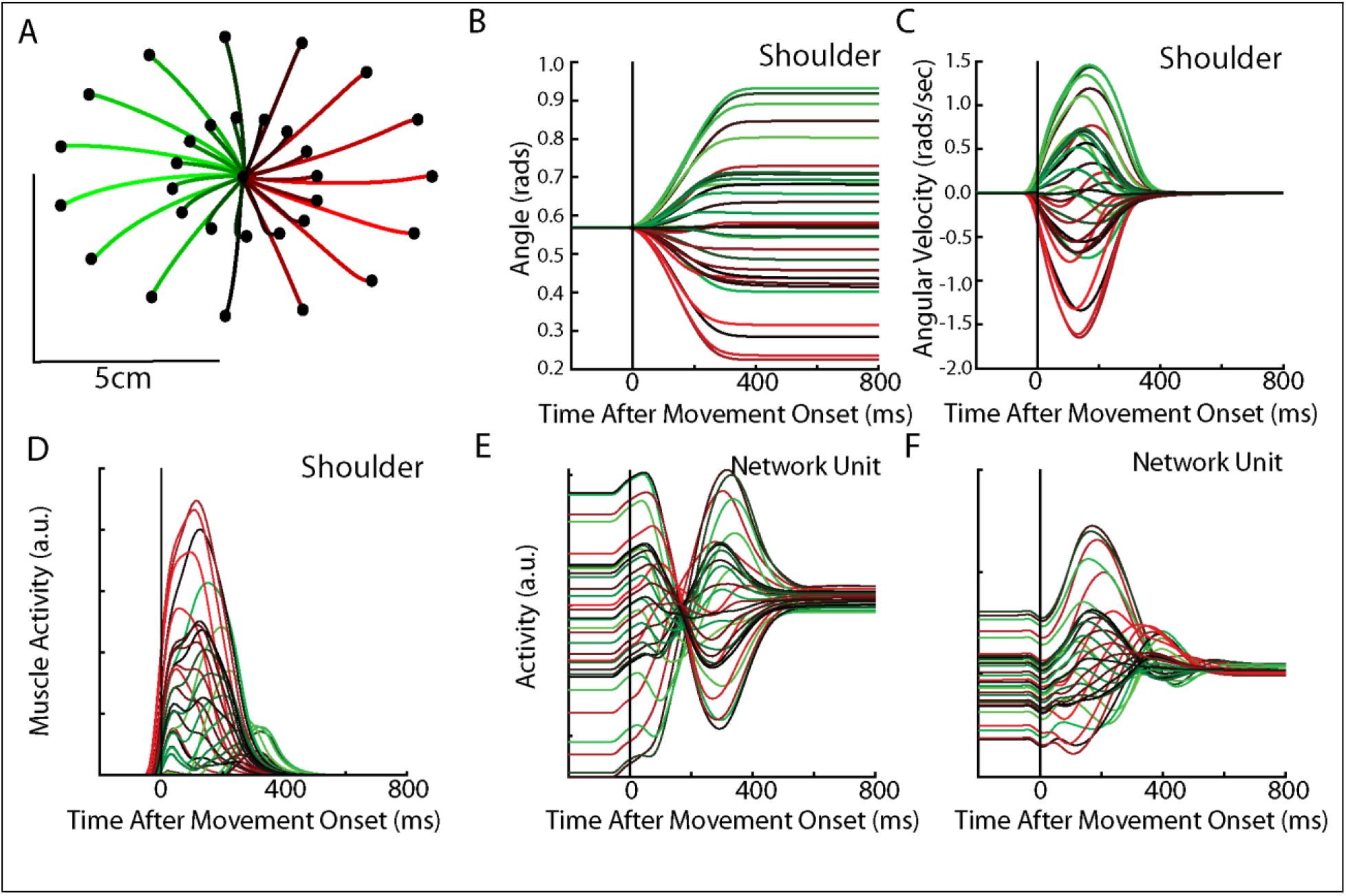
Delayed reach task by the network. A) The hand paths by the model’s arm from the starting position (center) to the different goal locations (black dots). Goals were placed 2cm and 5cm from the center location. B-C) Shoulder angle and angular velocity aligned to movement onset. D) Activity of the shoulder extensor aligned to Go cue onset. E-F) The activities of two example units from the output layer of the network.

We applied jPCA analysis to the output layer of the network and found clear rotational dynamics with rotational frequencies of 2.1Hz and 1.1Hz for the first and second planes, respectively (Figure 7A). These planes also captured 83% of the variance of the output-layer activity. When we examined the fit qualities, we found significant fits for the constrained and unconstrained dynamical systems with an R^2^ of 0.70 (p<0.001) and 0.83 (p<0.001), respectively (Figure 7B). Note, the ratio between the R^2^ for the constrained and unconstrained dynamical fits was 0.84, which is comparable to previous studies during reaching (Churchland et al., 2012) and indicate that the majority of the output layer’s dynamics displayed rotational dynamics.

**Figure 7.**
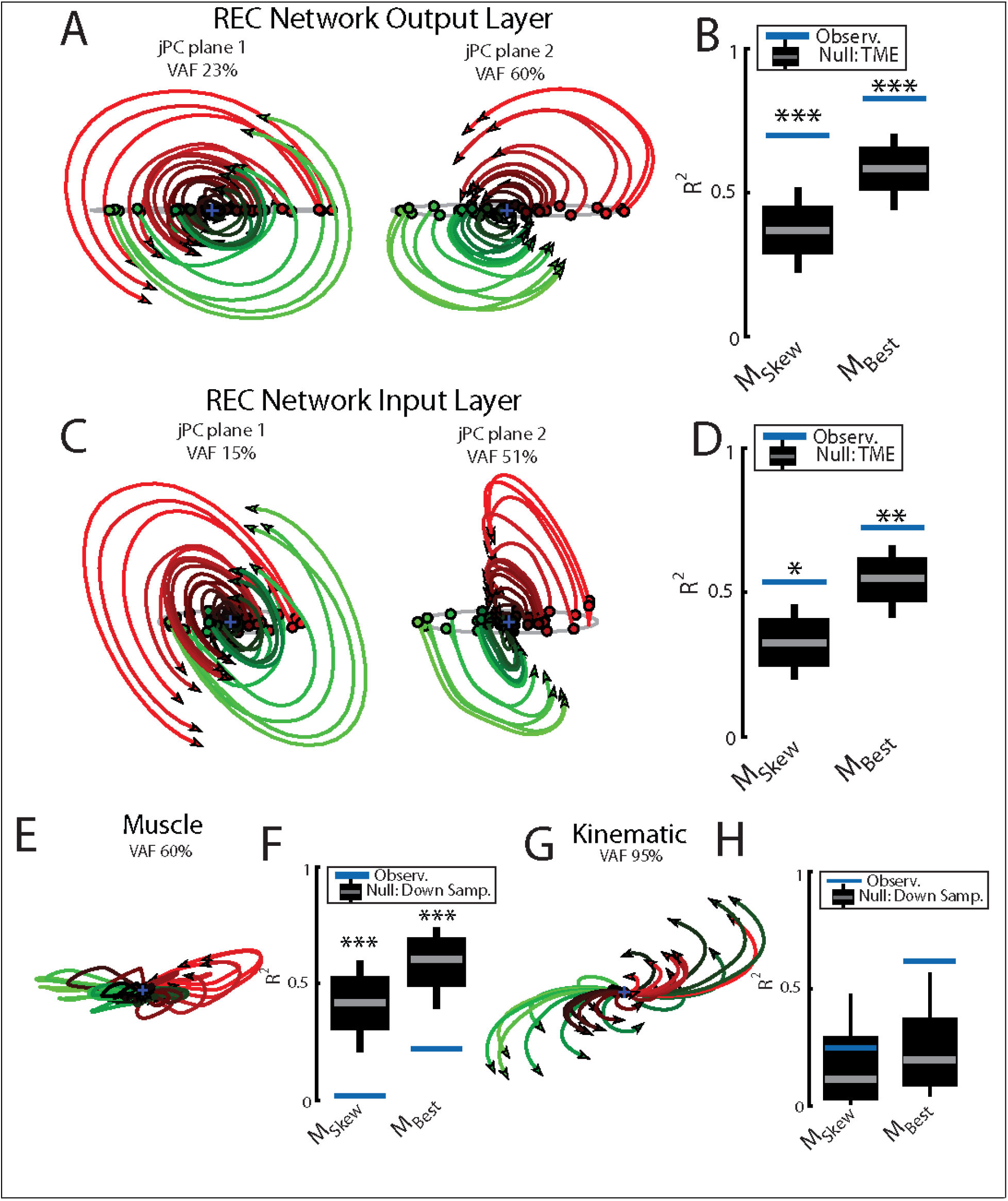
Population dynamics of the network during reaching. A) The top-2 jPC planes from the output layer of the network during reaching. B) Goodness of fits for the network activity to the constrained (M_Skew_ left) and unconstrained (M_Best_ right) dynamical systems. Null distributions were computed using tensor maximum entropy (TME). C-D) Same as A-B) for the input layer of the network. E-F) and G-H) Same as A-B) except for the muscle activities and kinematic inputs into the network, respectively. Null distributions were computed from the down-sampled neural activity. B, D, F, H) Grey bars denote the medians, the boxes denote the interquartile ranges and the whiskers denote the 10^th^ and 90^th^ percentiles. * p<0.05, **p<0.01, ***p<0.001.

We also examined the input layer of the network and found essentially the same results as the output layer (Figure 7C, D). Clear rotational dynamics were present rotating at 2.1 and 0.9 Hz in the top-2 planes, with significant fits for the constrained (R^2^=0.54, p=0.01) and unconstrained (R^2^=0.72, p=0.006) dynamical systems.

Next, we examined the dynamics of the muscle and kinematic signals. Similar to Churchland et al., (2012), we observed no rotational dynamics in the muscle activities (Figure 7E, F) and the fit for either dynamical system was significantly worse than the down-sampled network activity (constrained R^2^=0.02, p<0.001; unconstrained R^2^=0.24, p=0.01). In contrast, we observed rotational dynamics in the kinematic signals with a rotation frequency of 0.6Hz (Figure 7G, H). We found the kinematic signals were better fit by both dynamical systems and were comparable to the down-sampled neural activity (constrained R^2^=0.24 p=0.3; R^2^=0.62 p=0.06). Further, when we predicted the output layer’s activities using the combined sensory feedback (muscle, kinematics, GO cue, static inputs), we again found the predicted activities were highly similar (R^2^=0.99) to the output layer activities with virtually identical frequencies of rotation (Figure Supplementary 1B).

### Somatosensory cortex exhibits rotational dynamics while monkeys performed delayed reaching task

We explored if these dynamics were also present in somatosensory cortex during reaching, as previously observed in MC (Churchland et al., 2012). Monkeys H and C also completed a center-out reaching task using a manipulandum and data was recorded from area 2 (data from Chowdhury et al., 2020; Figure S6A). Note, these monkeys made slightly slower reaches (~400ms Figure S6B, C) than the reaches performed by the monkeys in Churchland et al., (2012) as well as our model simulations (both ~300ms).

We found clear rotational dynamics in area 2 with the top jPC plane having rotational frequencies of 1.0Hz and 1.7Hz for Monkeys H and C, respectively (Figure S6D). We also found significant fits for the constrained (Figure S6E, mean across monkeys R^2^=0.51, p<0.001 both monkeys) and unconstrained (R^2^=0.66, p<0.001) dynamical systems.

Next, examining the kinematics, we observed clear rotational dynamics in the top jPC plane with rotational frequencies of 1.3Hz and 1.2Hz for Monkeys H and C, respectively (Figure S6F). We also found significant fits for the constrained (Figure S6G, R^2^=0.39, Monkey H p<0.001, Monkey C p=0.02) and unconstrained (R^2^=0.51, Monkey H p<0.001, Monkey C p=0.01) dynamical systems.

### Neural networks without recurrent connections still exhibit rotational dynamics while performing posture and reaching tasks

Churchland et al., (2012) have suggested that these rotational dynamics emerge from the recurrent connections between neurons in MC. However, in our model, the sensory feedback into the network exhibited clear rotational dynamics that could contribute to the network’s dynamics. Thus, we explored if networks trained to perform the posture perturbation task without the recurrent connections (input and output layers) also exhibit rotational dynamics (i.e. 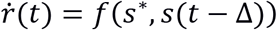)). We removed the recurrent connections in both the input and output layers of the network and optimized the network to perform the same posture task (NO-REC network). The network learned to bring the arm back to the central target when the external load was applied with similar kinematics as the REC network (data not shown).

Examining the output-layer activity, we still observed clear rotational dynamics with rotational frequencies of 1.0 and 0.74 Hz for the first and second planes, respectively (Figure 8A). These planes captured 92% of the variance of the network activity. When we examined the fit qualities, we found significant fits for the constrained dynamical system with an R^2^ of 0.43 (Figure 8B left; p=0.02), whereas for the unconstrained dynamical system we found a fit with an R^2^ of 0.54 but was not significant (Figure 8B right; p=0.3). As expected, output layer activities could be predicted from the sensory inputs with high accuracy (Figure S1C).

**Figure 8.**
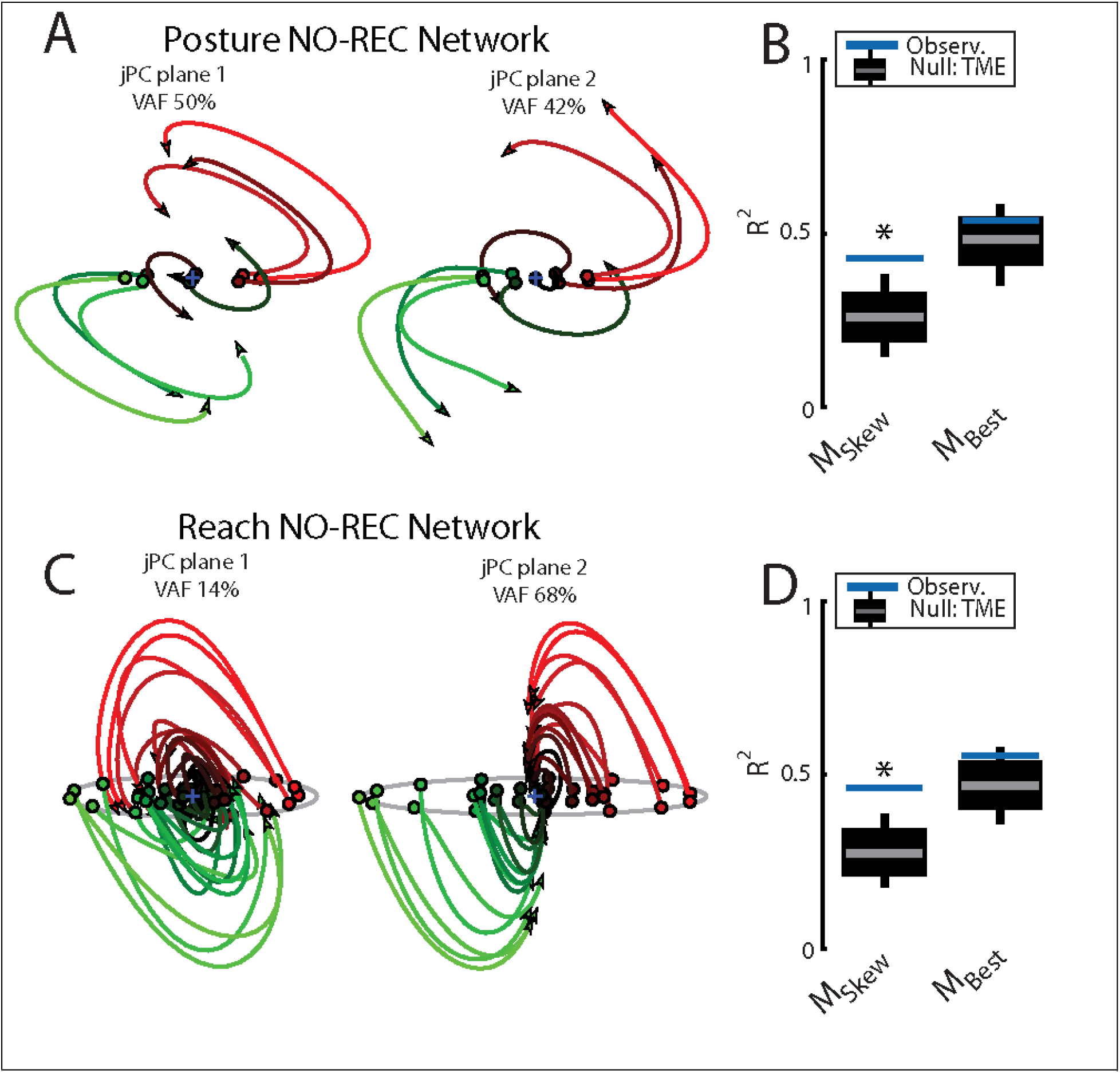
Population dynamics when trained without recurrent connections. Networks were trained to perform the posture and reaching tasks without the recurrent connections within the MC and input layers. A) The top-2 jPC planes from the output layer of the network during the posture task. B) Goodness of fits for the network activity to the constrained (M_Skew_ left) and unconstrained (M_Best_ right) dynamical systems. Null distributions were computed using tensor maximum entropy (TME). C-D) Same as A-B) for the output layer of the network during the reaching task. C, D) Grey bars denote the medians, the boxes denote the interquartile ranges and the whiskers denote the 10^th^ and 90^th^ percentiles. * p<0.05, **p<0.01.

Finally, we examined if the rotational dynamics would also occur in a network without recurrent connections for the center-out reaching task (NO-REC). We found this network exhibited good control of the limb with qualitatively similar hand paths to the targets as the REC network during reaching (data not shown). Examining the output layer’s dynamics, we observed rotational dynamics with rotational frequencies of 1.4 and 0.85Hz for the first and second planes, respectively (Figure 8C). These planes captured 82% of the variance of the network activity. When we examined the fit qualities, we found significant fits for the constrained dynamical system with an R^2^ of 0.46 (Figure 8D left; p=0.01), whereas for the unconstrained dynamical system we found a fit with an R^2^ of 0.56 but was not significant (Figure 8D right; p=0.15). Again, output layer activities could be predicted from the sensory inputs with high accuracy (Figure S1D).

## Discussion

The present study highlights how neural network models with sensory feedback and recurrent connections exhibit rotational dynamics in the network activities and in the sensory feedback from the limb, but not in muscle activities. These rotational dynamics were observed for a postural perturbation and a delayed reaching task, and critically, even without recurrent connections in the model. Similar tasks performed by monkeys also illustrate rotational dynamics not only in MC, but also in somatosensory areas and likely in sensory feedback signals related to joint motion. Thus, rotational dynamics are a characteristic that is present throughout the sensorimotor system, just not for muscles.

The standard equation to describe a linear dynamical system 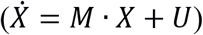 assumes the system evolves in time based on its own intrinsic dynamics (*M* ∙ *X*) and from inputs into the system (*U*) (Vyas et al., 2020). However, previous studies have argued that motor cortical dynamics are largely generated from intrinsic dynamics with inputs providing static information about the desired output and a nonselective GO cue to initiate movement (Churchland et al., 2012; Sussillo et al., 2015). This is supported by jPCA which fits neural activity using a linear dynamical system that only includes the term related to the intrinsic dynamics. This model captures rotational structure at the population level and can account for a substantial amount of neural variance. In contrast, limb muscle activity during reaching does not show these rotational dynamics. Furthermore, Sussillo and colleagues (2015) also found similar rotational dynamics in recurrent neural networks trained to generate the same patterns of muscle activity observed during reaching. Critically, these networks exhibited rotations despite only receiving relatively simple inputs (step function) and no sensory feedback. Thus, the dynamics were generated solely through recurrent connections in the model. Collectively, this leads to the interpretation that MC possesses a strongly interconnected network that generates patterns of muscle activity, and that this process is predominantly generated within MC.

The present study cannot directly refute that possibility, but it does provide several observations that clearly do not fit with this interpretation. Most critical is that our neural network model displayed rotational dynamics even when there were no recurrent connections and thus no intrinsic dynamics. Instead, rotational dynamics were generated by inputs to the network but could be inappropriately assessed as intrinsic dynamics. This suggests that rotational dynamics in MC may reflect internal dynamics, system inputs or any weighted combination of the two.

A second important observation is that we observed rotational dynamics in sensory feedback from the limb. Previous recurrent neural networks models of MC only used EMG-like signals for sensory feedback (Sussillo et al., 2015). However, primary and secondary afferents are critical sources of sensory feedback for limb control and their activity correlates with muscle length and change in that length (Cheney and Preston, 1976; Edin and Vallbo, 1990; Loeb, 1984). Our model and analysis of experimental data quantified joint angular position and velocity as a proxy of these sensory signals and found that they displayed rotational dynamics, similar to previous network models of control using kinematic variables (DeWolf et al., 2016; Susilaradeya et al., 2019). Furthermore, combined sensory feedback about kinematics and muscle activity could capture the high frequency rotations observed in the network activities indicating sensory feedback could provide rich dynamical signals for MC.

Another important observation in the present study is that rotational dynamics were observed not only in MC, but also in somatosensory cortex during the perturbation and reaching tasks. Rotational dynamics were observed in S1 (areas 3a and 1), A2 and A5, important components of frontoparietal circuits involved in the planning and execution of arm motor function (Chowdhury et al., 2020; Kalaska, 1996; Kalaska et al., 1990; Omrani et al., 2016; Takei et al., 2020). Thus, rotational dynamics are observed throughout frontoparietal circuits and likely in sensory feedback from the limb.

Although MC could still, in theory, generate the rotational dynamics exclusively through its recurrent connection, there are several reasons why inputs to MC are likely substantial during motor actions and contribute to its dynamics. Most notable is that behavioural level models of the motor system emphasize a dynamical systems perspective where various sources of information are rapidly processed to help guide and control ongoing motor actions. Optimal feedback control models have been influential as a normative model of voluntary control for almost 20 years (Scott, 2004; Todorov and Jordan, 2002). These types of controllers include two basic processes. First, state estimation where the present state of the body is optimally calculated from various sensory signals as well as from internal feedback generated using forward models. Second, a control policy uses this state estimate to generate motor commands to move the limb to a behavioural goal. These models predict many features of our motor system including that it is highly variable but also successful, and the ability to exploit redundancy while attaining a goal reflecting an interplay between kinematic errors and goal-directed corrections (Diedrichsen, 2007; Knill et al., 2011; Liu and Todorov, 2007; Nashed et al., 2012, 2014; Scott, 2016; Trommershäuser et al., 2005). A large body of literature highlights that goal-directed motor corrections to mechanical disturbances can occur in ~60ms and involve a transcortical pathway through MC (Matthews, 1991; Scott, 2004, 2012). These observations point to the importance of sensory feedback processing as a continuous rather than an intermittent process providing a continuous stream of input to brain circuits to guide and control motor actions (Crevecoeur and Kurtzer, 2018).

The dynamical systems view of MC activity developed from an attempt to understand the complex patterns of activity in M1, and how those dynamics lead to movement. This interpretation has tended to isolate processing by MC from the rest of the brain (but see Michaels et al., 2020) and that the objective of this processing is to generate patterns of muscle activity. However, this interpretation does not predict or explain behaviour – such as what the constraints or optimality criteria are that shape behavior or what computational problem the brain is trying to solve? These are exactly the problems addressed by optimal control models. Optimal control theory focusses on the importance of the entire circuit including sensory feedback for goal-directed control and has good explanatory power at the level of behaviour. Critically, it is the behavioural goal that is the fundamental objective as muscle activity can vary from trial-to-trial reflecting necessary corrective responses to deal with noise and errors. However, optimal control theory will need additional assumptions and structure to explain the nature of neural processing. Thus, the two classes of models have the potential to be complementary and work together.

One feature not captured by our model is that complex multi-phasic activity patterns precede movement onset by 100-150ms, and this observation has been used as evidence for autonomous MC dynamics (Churchland et al., 2012; Schroeder et al., 2019; Sussillo et al., 2015). Obviously sensory feedback of the movement cannot play a role in generating these early responses, which must instead occur through internal processing, including inputs from other brain regions (Sauerbrei et al., 2020). Though of course these inputs can include sensory feedback about the state of the limb and the movement goal (Ahmadi-Pajouh et al., 2012; Ames et al., 2019; Pruszynski et al., 2008, 2014). In any case, even if the pre-movement dynamics in MC were autonomous, this would not imply that MC continues to behave as an autonomous system during movement. Instead, our results show that sensory feedback is likely to contribute heavily to MC dynamics during movement.

How inputs conveying sensory and internal feedback are processed by MC remains an important and poorly understood problem in motor control. Recent studies have suggested that MC uses an initial planning stage when processing visual feedback during movement. Stavisky et al., (2017) showed that the initial visual feedback response to a shift in hand position during reaching may be transiently isolated from the activity associated with generating motor output. However, as we show here, this latter activity may still reflect sensory and internal feedback. Similarly, Ames et al., (2019) showed that jumping the location of the goal during reaching to a new location generated activity patterns that were similar to the patterns generated when planning a separate reach to the new goal’s location. This planning stage may reflect an update to the control policy given the visual error, resembling model predictive control (Dimitriou et al., 2013) and it remains an open question if these feedback responses to systematic errors (visual shift or mechanical load) evoke the same activity patterns in MC as motor noise (Crevecoeur et al., 2012).

However, new techniques will be required to better explore how inputs are processed by MC. Recent methods that exploit simultaneous recordings from multiple brain areas provide a promising tool to identify input signals to a given circuit (Kohn et al., 2020; Perich et al., 2018; Semedo et al., 2019). Using these techniques, Perich et al., (2020) provides evidence of a communication subspace between the somatosensory and motor cortices that contributes to a substantial amount of the variance in MC, consistent with inputs playing a key role in motor cortical dynamics. Studies that perturb neural circuits through stimulation, cooling probes or optogenetics will also provide valuable insight into how inputs are transformed by MC (Guo et al., 2020; Hore et al., 1977; Li et al., 2016; Nashef et al., 2018, 2019; Perich et al., 2020; Svoboda and Li, 2018; Takei et al., 2020). For example, deactivating cerebellar output can substantially impact preparatory activity in the MC and feedback responses to mechanical loads (Chabrol et al., 2019; Conrad et al., 1974; Gao et al., 2018; Meyer-Lohmann et al., 1975). Sauerbrei et al., (2020) also recently uncovered how the sudden loss of input from motor thalamus results in a collapse of motor cortical dynamics. These techniques and future advancements will be needed to tease apart dynamics generated internally versus dynamics generated from external sources.

## Acknowledgements

We thank Kim Moore and Helen Bretzke for their laboratory and technical assistance. This work was supported by grants from the Canadian Institute of Health Research. KPC was supported by an Ontario Graduate Scholarship. SHS was supported by a GSK chair in Neuroscience.

## Author Contributions

Conceptualization, H.T.K., K.P.C., T.P.L, M.O., and S.H.S; Methodology, H.T.K., K.P.C., and S.H.S; Writing, H.T.K., K.P.C., T.L.P, P.N.S., and S.H.S.; Formal Analysis, Investigation, H.T.K., K.P.C.; Funding Acquisition, E.F. and S.H.S.; Supervision, E.P and S.H.S.

---

**Supplementary Figure 1.**
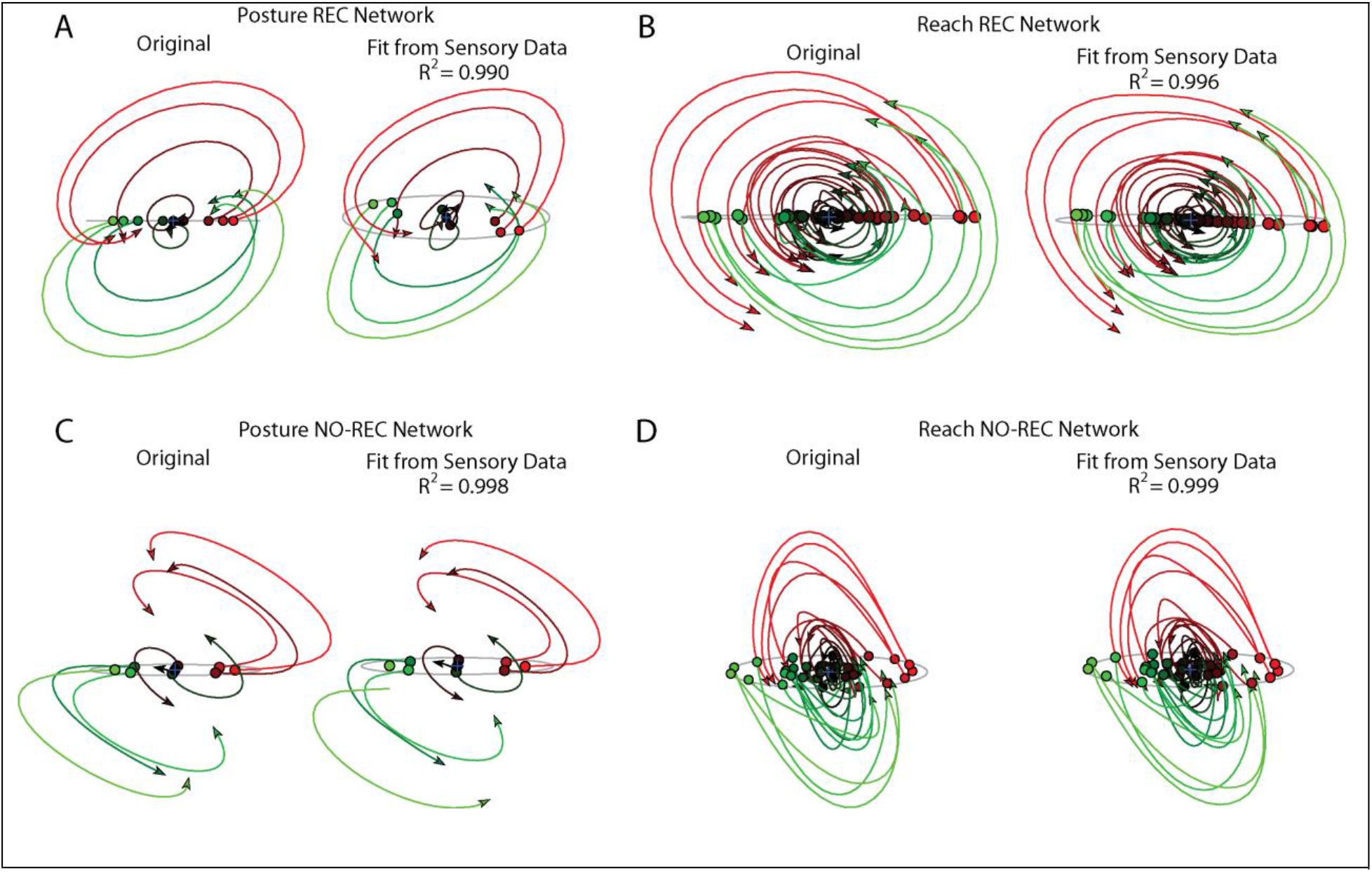
Predicting output layer trajectories using sensory input. A) The top jPC plane from the output layer activities (left) and the predicted activity using only sensory feedback (right) during the perturbation posture task. R^2^ reflects the fit quality across all 6 jPC planes. B) Same as A) for the center-out reaching task. C-D) Same as A-B) except for the NO-REC networks.

**Supplementary Figure 2.**
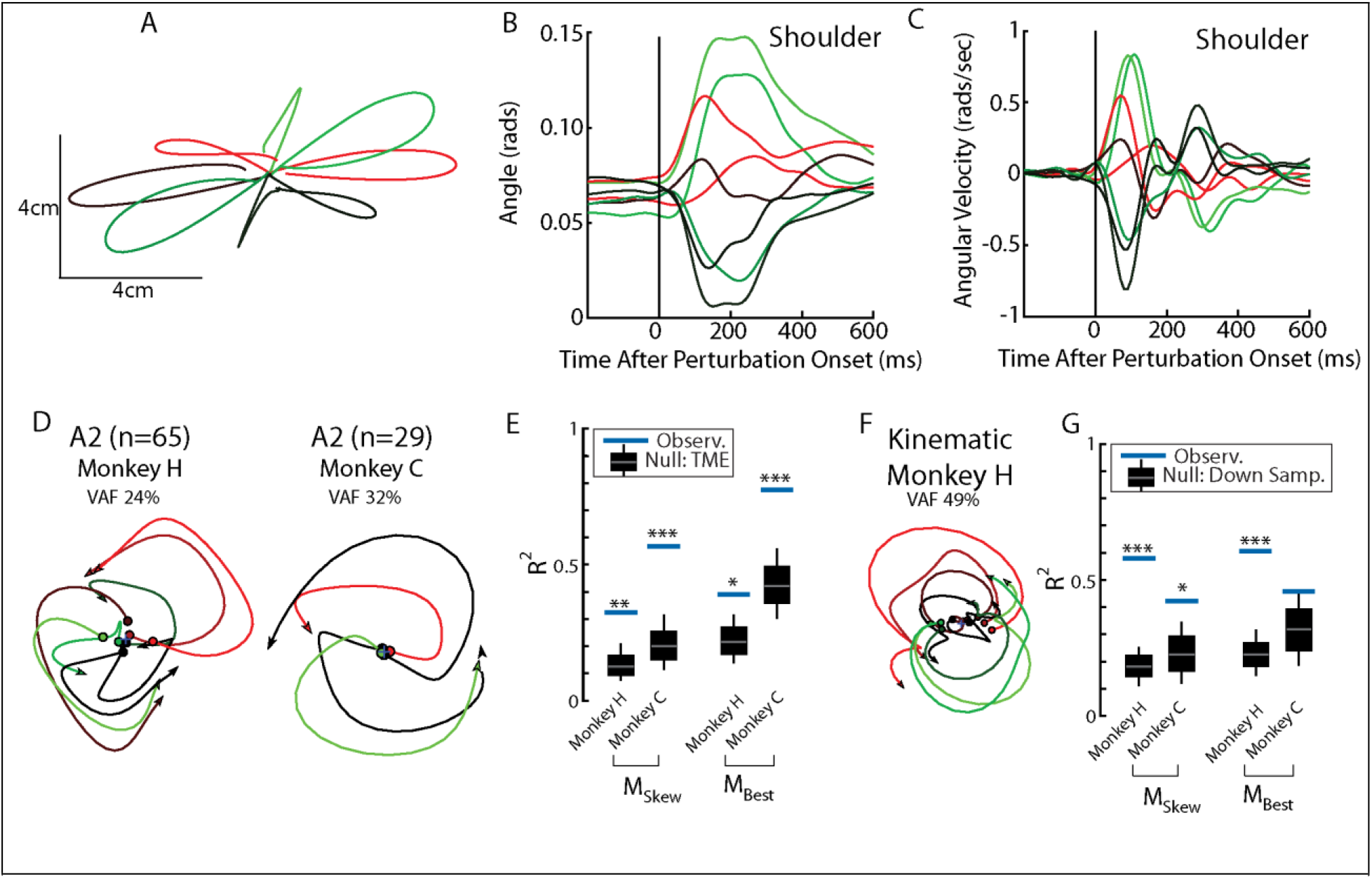
Population dynamics in somatosensory cortex during posture task from Chowdhury et al., (2020). A) Hand paths for Monkey H using an endpoint manipulandum where loads were applied that displaced the hand from the starting position. B, C) The shoulder flexion angle and angular velocity across the load directions. D) The top jPC plane from activity recorded in somatosensory area 2. E) Goodness of fits to the constrained (M_Skew_ left) and unconstrained (M_Best_ right) dynamical systems. Null distributions were computed using tensor maximum entropy. F-G) Same as D-E) except for the kinematic signals. Null distributions were computed from the down-sampled neural activity. Data from Chowdhury et al., (2020). * p<0.05, ** p<0.01, *** p<0.001.

**Supplementary Figure 3.**
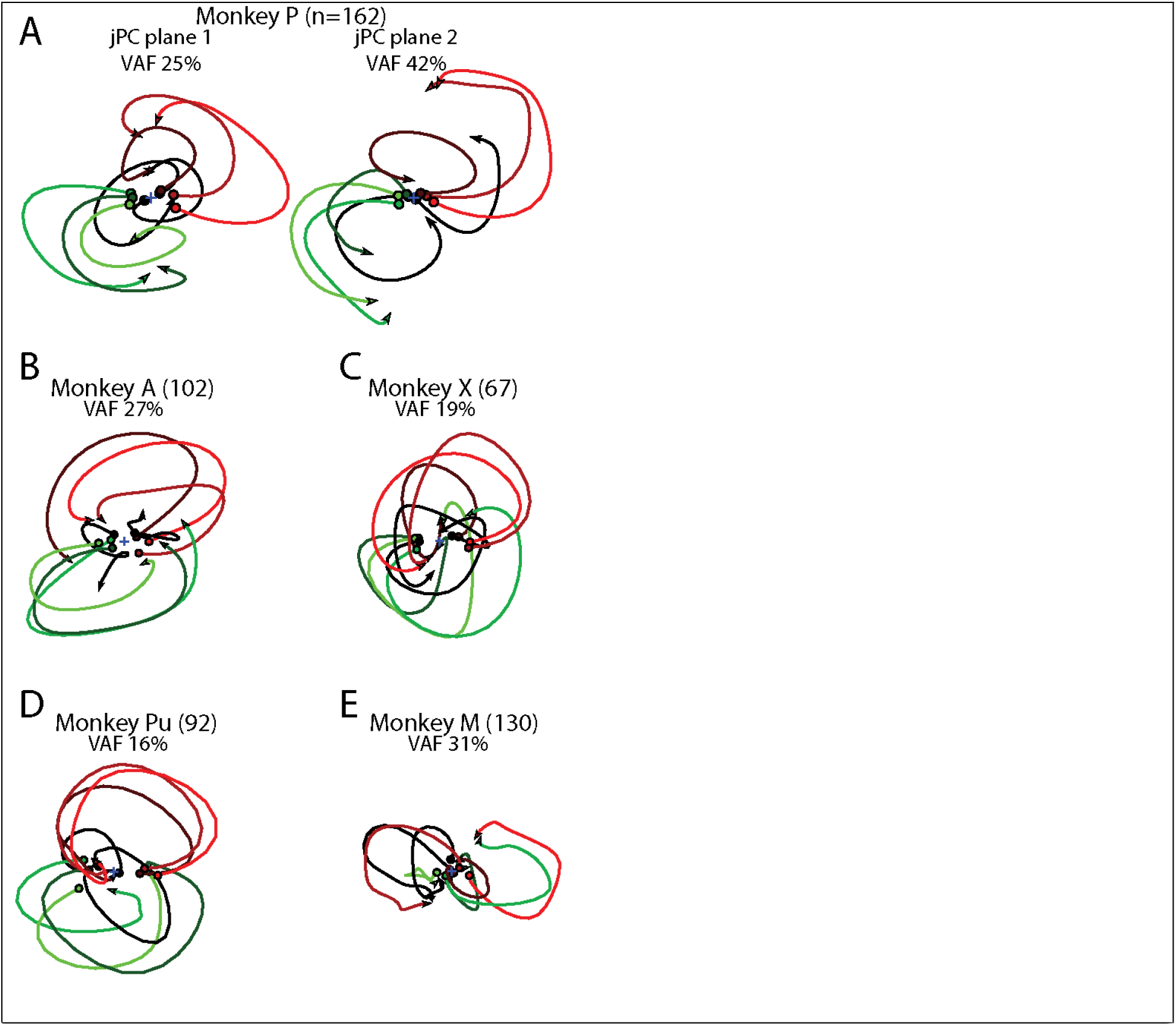
Population dynamics in motor cortex for individual monkeys. A) The top-2 jPC planes from activity recorded in motor cortex in Monkey P. B) The top jPC plane from activity recorded in motor cortex in Monkey A. C-E) Same as B) for Monkeys X, Pu, and M.

**Supplementary Figure 4.**
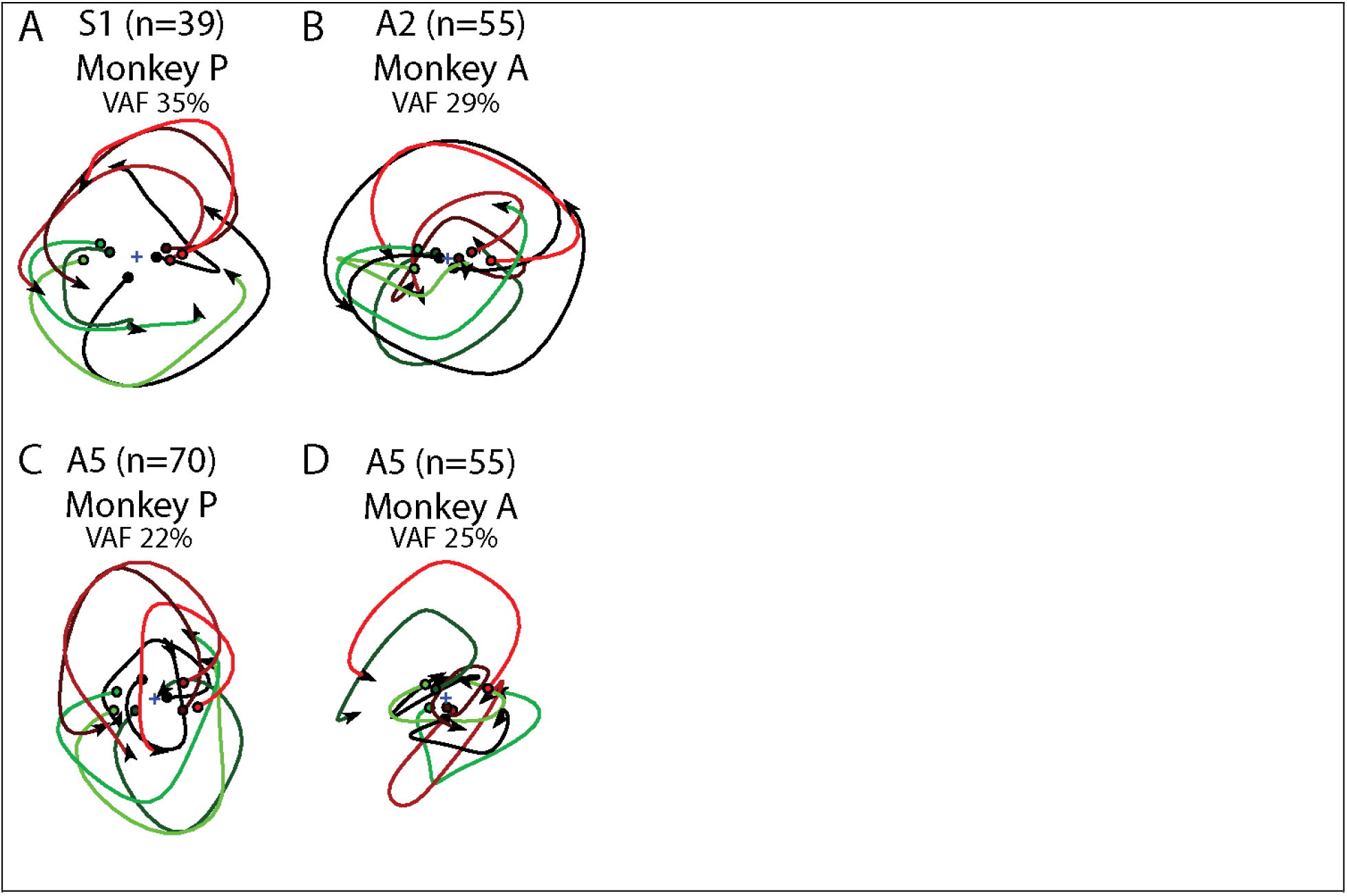
Population dynamics in somatosensory cortex for individual monkeys. Data are presented the same as Supplementary Figure 3 for S1 in Monkey P (A), A2 in Monkey A (B), A5 in Monkey P (C) and A5 in Monkey A (D).

**Supplementary Figure 5.**
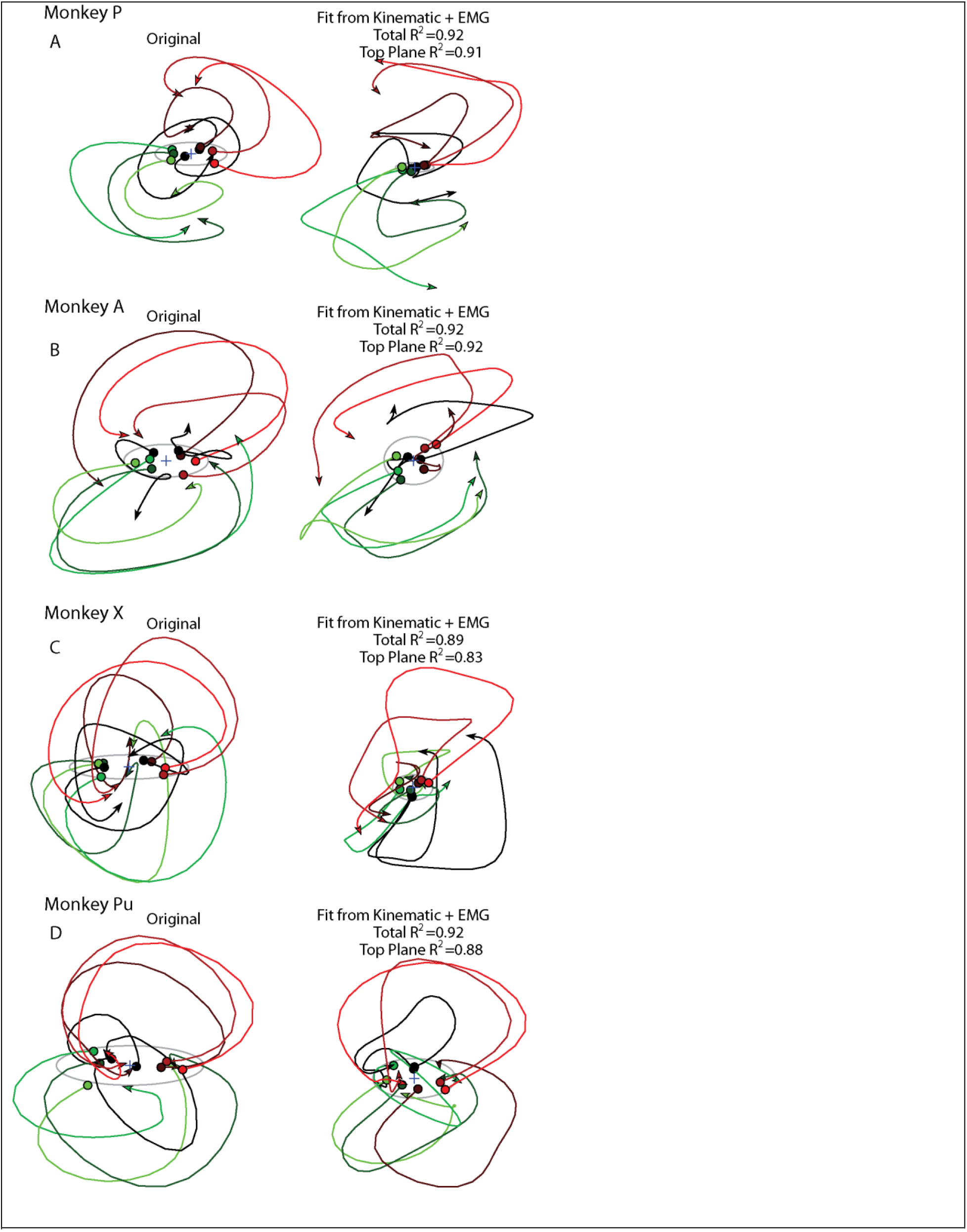
Predicting M1 activity using kinematic and muscle activities. Data presented the same as in Figure Supplementary 1 except for individual monkeys.

**Supplementary Figure 6.**
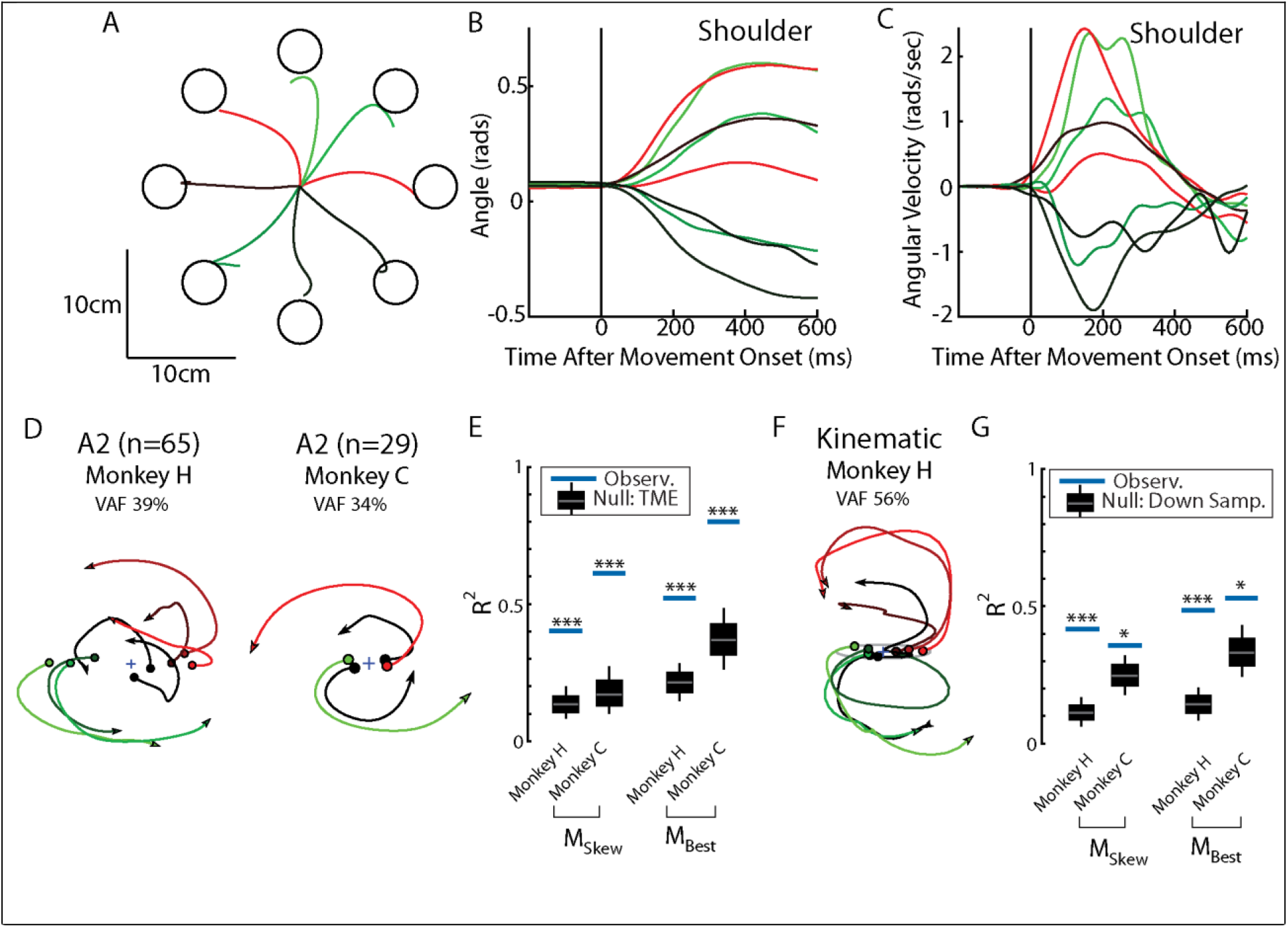
Population dynamics in somatosensory cortex during reaching from Chowdhury et al., (2020). A) Hand paths for Monkey H using an endpoint manipulandum to reach to different targets located in a center-out pattern. Targets were placed 12cm from the starting position. B-C) The shoulder flexion angle and angular velocity across the different reach directions (shoulder flexion angle defined in Chan and Moran, 2006). D) The top jPC plane from activity recorded in somatosensory area 2. E) Goodness of fits to the constrained (M_Skew_ left) and unconstrained (M_Best_ right) dynamical systems. Null distributions were computed using tensor maximum entropy (TME). F-G) Same as D-E) except for the kinematic signals. Null distributions were computed from the down-sampled neural activity. * p<0.05, ***p<0.001.

## Materials and Methods

### Two-link arm model

We constructed a two-link model of the upper arm as detailed in Lillicrap and Scott, (2013). The model was constrained to move in a horizontal two-dimensional plane and incorporated arm geometry and inter-segmental dynamics. The dynamics of the limb were governed by

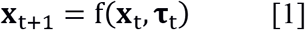

Where, ‘**x**_t_’ is the vector state of the arm at time ‘t’ and was composed of the angular positions and velocities of the elbow and shoulder joints [θ_elb_, θ_sho_, 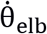, 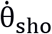]. ‘**τ**_t_’ is the two-dimensional vector of torques applied to the shoulder and elbow joints at time ‘t’. We incorporated 6-lumped muscle actuators that moved the arm, which included 4 mono-articular and 2 bi-articular muscles. These muscles received input from the neural network and exhibited force-length and force-velocity dependent activation properties (Brown et al., 1999). Muscle forces (m_t_) were converted to joint torques by computing the product between each muscle’s force output with their respective moment arm. The parameters for the arm dynamics, moment-arm matrix and the muscle force-length/velocity (F-L/V) properties were drawn from the literature (Brown et al., 1999; Cheng et al., 2000; Graham and Scott, 2003). The continuous arm dynamics were discretized and solved using Euler’s integration with a time step (dt) of 10ms.

### Network description

We used a recurrent neural network (RNN) composed of two layers to control the arm model. Both layers had recurrent connections between units within each layer and all units had leaky-integration properties and a standard sigmoid activation function.

The first layer received inputs (**s**_t_) composed of a step signal representing the desired joint state 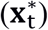, delayed (Δ=50ms) state feedback from the arm (**x**_t−Δ_, joint angles and angular velocities) and delayed muscle activations (**m**_t−Δ_). For the reaching task we also included a condition-independent binary ‘GO’ cue to indicate when the network should initiate movement. This signal was applied as a step function smoothed with a 20ms s.d. Gaussian kernel (high indicates hold command, low indicates move command). The dynamics of the first layer (referred to as input layer) were governed by

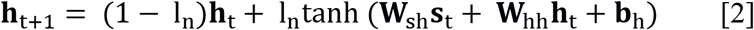

Where, **h**_t_ is the vector of unit activities for the input layer, ‘l_n_’ is the ratio between the simulation time-step (dt) and the time-constant of the network units (τ_n_), hence l_n_ = dt/τ_n_. **W**_sh_ is the weight matrix that maps the inputs to the activities of the input layer, **W**_hh_ is the weight matrix for the recurrent connections between units in the input layer, and **b**_h_ is the bias (or baseline) for the first layer activities.

The second layer (output layer) received input from the input layer and its dynamics were governed by

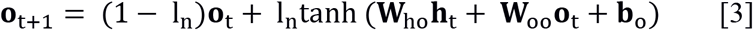

Where, **o**_t+1_is the vector of unit activities for the output layer, **W**_ho_ is the weight matrix that maps the input layer activities to the output layer activities, **W**_oo_ is the weight matrix for the recurrent connections between units in the output layer, and **b**_o_ is the bias (or baseline) for the outputlayer activities.

The output layer provides inputs to the 6 muscles used to control the limb. The muscle activities (**m**_t_) were governed by,

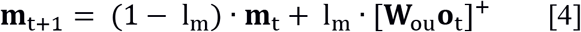

**W**_ou_ is the weight matrix that maps the activities in the output layer to the lumped muscle actuators, and l_m_ is the leak time constant for the muscle given by, l_m_ = dt/τ_m_.

We also examined networks where we removed the recurrent connections from each layer by effectively setting **W**_hh_, **W**_oo_ to zero for the entire simulation and optimization (NO-REC networks).

For all simulations, the input and output layers were composed of N = 500 units each and the time constants of network units (τ_n_) and muscle units (τ_m_) were 20ms and 50ms, respectively. The weight matrices were initialized from a gaussian distribution centered on zero with a standard deviation of ± 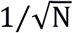. All the bias vectors [**b**_h_, **b**_o_] were initialized to 0.

### Choice of sensory inputs into network

Our model receives delayed sensory feedback from the periphery composed of the angles and angular velocities of the joints as well as the muscle activities. We think these are reasonable inputs into the network based on known properties of proprioceptors. Activity of muscle spindles are known to signal muscle length and velocity (Cheney and Preston, 1976; Edin and Vallbo, 1990; Loeb, 1984), which could be used to form an estimate of joint angle and angular velocity (Scott and Loeb, 1994). Activity of Golgi tendon organs signal muscle tension and correlate with muscle activity (Houk and Henneman, 1967; Nichols, 2017; Prochazka and Wand, 1980).

### Task descriptions

We trained the network to perform a posture perturbation task similar to our previous studies (Heming et al., 2019; Omrani et al., 2014; Pruszynski et al., 2014). The network was required to keep the arm at a desired position while the limb was displaced by loads applied to the shoulder and elbow joints. Eight torques (of magnitude 0.2Nm) were used consisting of elbow flexion (EF), elbow extension (EE), shoulder flexion (SF), shoulder extension (SE), and the four multi-joint torques (SF+EF, SF+EE, SE+EF, SE+EE). Importantly the network did not receive any explicit information on the direction of the applied load and has to use the delayed sensory feedback to produce appropriate compensation.

We also trained separate instances of the network to perform a delayed center-out reach task that required the network to hold the arm at a starting position for 500ms. Afterwards, a GO cue appeared signaling the network to move to the target within 500ms. We had the network reach to 32 different targets spaced radially around the starting position with half of the targets located 2cm away from the starting position, and the remaining half were placed 5cm away from the starting position. The network then had to hold at the reach target for the remainder of the trial (~500ms). Note, for our simulations we used a fixed time delay (represented by the GO signal) for when the network should initiate a reach to decrease optimization time. Simulations with a variable delay yielded virtually the same results.

### Network optimization

For optimizing the networks, we defined the loss function (‘l’) over a given trial (i) as

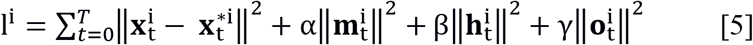

Where *α*, *β*, *γ* are penalization weights. The first term of the loss function is the vector norm between the desired limb kinematic state 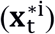 and the current limb kinematic state 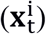. The second term penalizes the total muscle activity, and the third and fourth terms penalize high network activities for the first and second layers, respectively.

In the posture perturbation task, the desired limb state was static irrespective of the direction of external torques, and the kinematic term considered the norm of the difference between the desired state of the arm and the actual state 1000ms after the time of load application. In the reach task, the desired limb state was defined as the location of the reach target on that trial and the kinematic error was penalized 500ms after the GO cue was presented. Similar to the posture task, the muscle and network activities were penalized during the entire reach task.

The network parameters were determined by minimizing the total cost ‘J’ from summing the individual trial loss functions across different movement types (i.e. the 9 load combinations in the posture task or 32 target locations in the reach task).

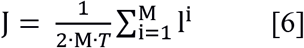

We optimized the network by applying back-propagation through time (Werbos, 1990). This requires us to compute the cost-gradient 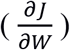 with respect to the adjustable network parameters ***W*** = [***W***_*sh*_, ***W***_*hh*_, ***W***_*ho*_, ***W***_*oo*_, ***W***_*ou*_, ***b***_*h*_, ***b***_*o*_]. Since, the total cost depends upon the kinematic state of the arm (***x***_*t*_), the optimization problem involves calculating the Jacobian of the arm dynamics 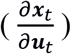 at each time-step, as presented in Stroeve, (1998). Our simulations were implemented in Python and PyTorch machine learning library (Paszke et al., 2017). Optimization was performed using the Adam algorithm (Kingma and Ba, 2017) and performed until the network generated successful limb trajectories and the error had decreased to a small, constant valuer (approx. 1e-4) for at least 500 epochs. For all the simulations, the hyper-parameters were fixed at *α*=1e-4/1e-3, *β* = 1e-5/1e-6 and *γ* = 1e-5/1e-6; although comparable network solutions were obtained for a broad range of these hyper-parameter values. Note, in the posture task, during a delayed period before the application of any load, the muscle activities were penalized with a higher *α* = 1e-2 to ensure that the muscles were not active by default at a higher baseline to counter-act the upcoming load.

### Neural recordings

We analyzed neural activity from fronto-parietal areas when monkeys performed a posture perturbation task that had been previously collected (Chowdhury et al., 2020; Heming et al., 2019; Omrani et al., 2014, 2016; Pruszynski et al., 2014). Briefly, Monkeys P, A, X, Pu, and M had their arms placed in a robotic exoskeleton that restricted the animal’s movements to motion of the shoulder and elbow joints in a 2-d horizontal plane. These animals performed almost the exact same posture perturbation task as the network. However, different load magnitudes were used for each monkey depending on their physical capabilities (Monkeys P, X =0.2Nm, A=0.4Nm, Pu=0.2Nm, M=0.34Nm). Also, for some recordings in Monkey P, X and M the load was removed 300ms after it was applied. Given that we were interested in the earliest feedback response, we included these recordings. Data for Monkeys H and C were from Chowdhury et al., (2020) where the monkeys performed a similar task using a robotic manipulandum and where 2N forces were applied to the manipulandum that lasted 125ms (London and Miller, 2012).

Monkeys H and C also performed a delayed center-out reaching task (Chowdhury et al., 2020; London and Miller, 2012). Goal targets were arranged radially around the starting position at a distance of 12.5cm. For Monkeys H and C, eight and four different goal locations were used, respectively. After the delay period, the monkeys had to reach for the goal location within ~2seconds for a successful reach.

Single tungsten electrodes were used to record cortical activity from Monkeys P, A and X and floating micro-electrode arrays were used to record from Monkeys M, Pu, H and C. Primary motor cortex activity was recorded from Monkeys P, A, X, Pu and M. Premotor cortex activity was also recorded from Monkeys P and A, which were pooled with the primary motor cortex neurons. Primary somatosensory area 1 (areas 3a and 1) and parietal area 5 were recorded from Monkey P. Primary somatosensory area 2 and parietal area 5 were recorded from Monkey A. Primary somatosensory area 2 was recorded from Monkeys H and C.

Spike timestamps were convolved with a gaussian kernel with a standard deviation of 30ms. For displaying the single neuron responses only, timestamps were convolved with a half-gaussian kernel (SD 30ms) that only estimated the instantaneous firing rate using spikes from the past. This prevented the appearance during the posture perturbation task that changes in firing rates preceded the onset of the load.

### Muscle recordings

Muscle activity was recorded percutaneously by inserting two single-stranded wires into the muscle belly (Scott and Kalaska, 1997). Stimulation was used to confirm the penetrated muscles. We recorded from the main extensor and flexor muscles of the shoulder and elbow including triceps (lateral and long), biceps (long and short), deltoids (anterior, medial and posterior heads), brachioradialis, supraspinatus and pectoralis major. From each monkey we recorded a subset of these muscles that included a mixture of flexor and extensor muscles for both the shoulder and elbow joints.

### jPCA analysis

We performed jPCA analysis on the neural network similar to Churchland et al., (2012) using code available at https://churchland.zuckermaninstitute.columbia.edu/content/code. We constructed matrices X that contained the activities of all neurons in the network for every time point and condition (i.e. load combinations or reach directions). These matrices had NxCT dimensions, where N is the number of neurons in the network, C is the number of conditions, and T is the number of timepoints. The mean signal across conditions was subtracted at each time point and activity was soft normalized by the activity range plus a small constant (5e-4).

Principle components analysis (PCA) was applied to X and the top-6 principle components were used to reduce X to *X*_*Red*_ (6xCT dimensions). We numerically calculated the derivative of *X*_*Red*_ yielding 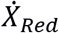, and fit a linear dynamical model which found a relationship between *X*_*Red*_ and 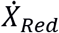

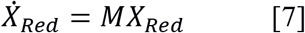

Where M is a 6×6 weight matrix. We assessed the model’s fit by calculating the coefficient of determination (*R*^2^).

With no constraint on M, any linear dynamical system could be captured by this equation including oscillators, point and line attractors, etc. We compared how an unconstrained M performed with a fit where we constrained M to be skew symmetric (*M*_*Skew*_). This restricted the possible dynamical systems to systems with oscillatory dynamics. Skew-symmetric matrices have pairs of eigenvectors with eigenvalues that are complex conjugates of each other. These eigenvector pairs were found from *M*_*Skew*_ and the corresponding activity generated 2-dimensional jPCA planes. *M*_*Skew*_ generates 3 jPCA planes and the planes were ranked by their eigenvalues (i.e. the speed of the rotational dynamics) from highest to lowest. The amount of variance each plane captured of the original matrix X (VAF) was calculated and normalized by the total amount of variance in the original matrix X.

jPCA analysis was also applied to the kinematic feedback signals from the plant (normalization constant 0), the muscle activity produced by the network (0), the recorded neural activity (5sp/s) and the recorded EMG activity (0). Since there are fewer kinematic and muscle signals than neural signals, we only examined activity in the top-two kinematic components, and the top-four muscle components. For the posture task, jPCA analysis was applied for the first 300ms after the load onset for the neural recordings. For the network, jPCA analysis was applied from 70-370ms after the load onset to reflect the 50ms delay in sensory feedback processing. Similar results were obtained using 0-300ms epoch. For the reaching data, jPCA analysis was applied for the first 300ms after the start of movement.

### Tensor maximum entropy

We tested our findings against the hypothesis that rotational dynamics are a byproduct of the tuning and smoothness properties of neurons. We employed tensor maximum entropy to generate surrogate datasets (Elsayed and Cunningham, 2017) using code available at https://github.com/gamaleldin/TME. This method generates surrogate data sets that preserve the covariances across neurons, conditions and time but not their interactions as required for rotational dynamics. Surrogate data sets were then sampled from this distribution and the jPCA analysis was applied to each data set (1000 iterations).

### Down-sampling neuron activity

For the muscle and kinematics, assessing whether the observed rotational dynamics were significant or not was complicated by the fact that there were fewer muscle and kinematics signals. Indeed, neural population dynamics deemed significant using TME were no longer significant after down sampling the neural population to match the number of kinematic and muscle samples. Instead, we assessed whether the rotational dynamics in the muscle or kinematic signals were more dynamical than neural activity after correcting for the number of signals. We randomly sampled neurons from the neural population to match the number of muscles or kinematic signals and applied jPCA analysis to the resulting population activity. This was repeated 1000 times.

